# Pan-cancer analysis reveals tumor microbiome associations with host molecular aberrations

**DOI:** 10.1101/2023.04.13.536730

**Authors:** Chenchen Ma, Changxing Su, Jiaxuan Li, Jiuxin Qu, Shimin Shuai

## Abstract

Host-microbiome interaction is known to play a pivotal role in the cancer ecosystem, yet the associations have not been systematically investigated at the pan-cancer and the multi-omics level. Here, we evaluated nearly 10,000 samples across 32 cancer types collected from The Cancer Genome Atlas (TCGA), to investigate the association between the tumor microbiome (taxa, n=1,630) and tumor microenvironment composition (cell types, n=20), epigenome (CpG island methylation, n=30,716), transcriptome (gene expression, n=10,216) and proteome (protein expression, n=193). We identified 836,738 candidate associations between the tumor microbiome and host molecular aberrations across multiple cancers. Besides cancer-specific associations, we also revealed recurrent pan-cancer associations between microbes (*Lachnoclostridium*, *Flammeovirga*, *Terrabacter* and *Campylobacter*) and immune cells, as well as between microbes (*Collimonas* and *Sutterella*) and fibroblasts, which were further validated by cell type estimations derived from pathological images and methylation data. We also identified several potential microbe and gene/protein expression associations mediated by DNA methylation using the sequential mediation analysis. Furthermore, our survival analysis demonstrated that tumor microbes may affect the patient’s overall survival and progression-free survival. Finally, a user-friendly web portal, Multi-Omics and Microbiome Associations in Cancer (MOMAC) was constructed for users to explore potential host-microbe interactions in cancer.

**Graphical abstract:** 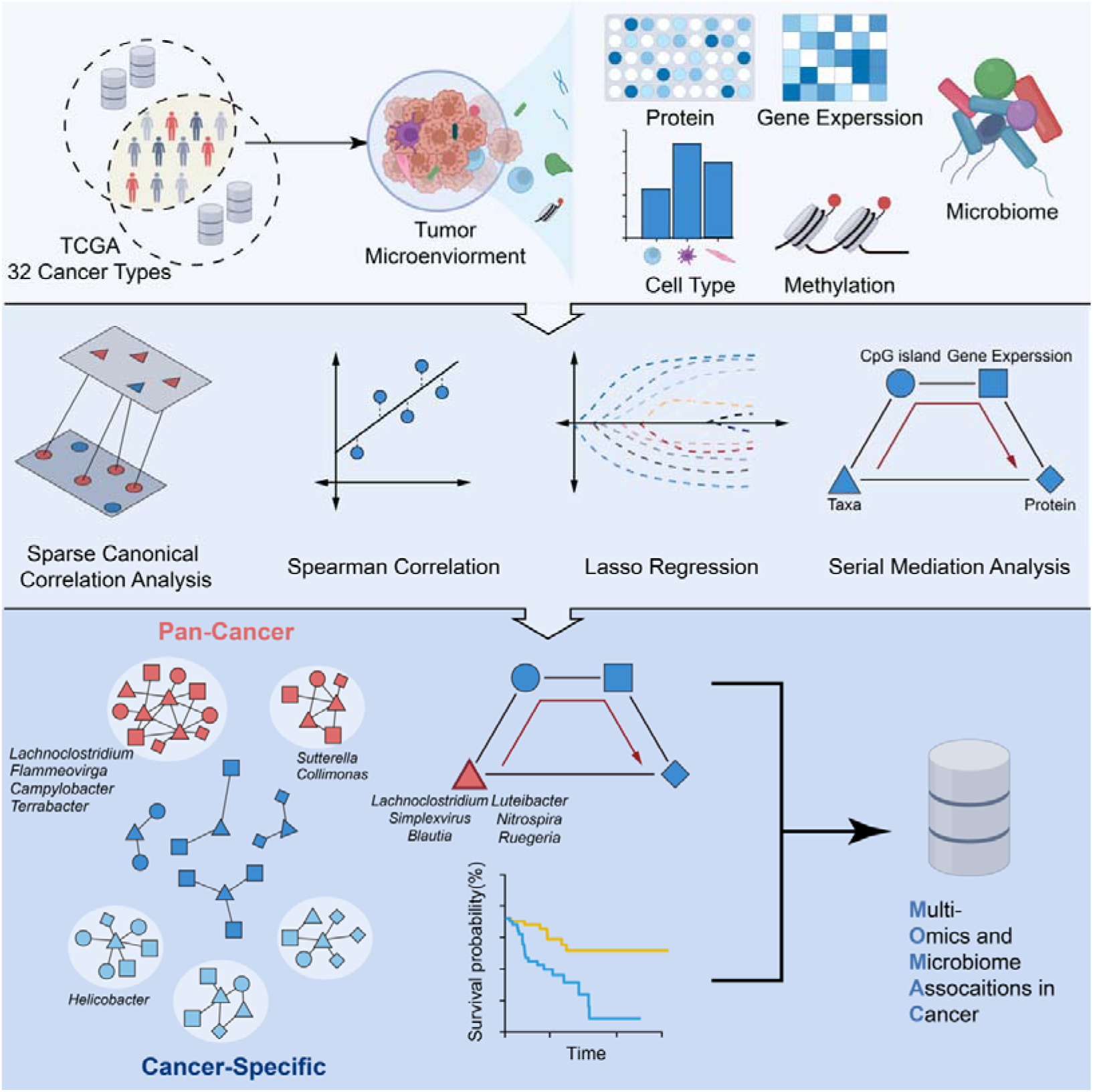

## Introduction

Recent large-scale analyses reveal that tumor-resident bacteria, archaea, viruses and fungi, collectively called the tumor microbiome ^1–4^, are a key component of the cancer ecosystem as “permanent resident” rather than “short-term tenant” ^5^. Several functional and association studies have revealed that the tumor microbiome may play crucial roles across the entire lifespan of cancer, such as promoting oncogenesis ^6^ and metastasis ^7^, as well as modulating the cancer ecosystem and treatment response ^8, 9^.

Treatment efficacy and cancer development may be significantly impacted by the interactions between the tumor microbiome, tumor cells, and immune cells in the cancer ecosystem ^9, 10^, which is largely orchestrated by inflammatory cells ^11^, but little is known about the precise action modes of the interplay between the tumor microbiome and host cells in the cancer ecosystem. One hypothesis could be that the tumor microbiome contributes to tumor progression and development via affecting the host immune responses and cancer-related signaling pathways ^4^. However, systematic analysis of associations between the tumor microbiome and host molecular profiles such as DNA methylation, RNA expression and protein levels in pan-cancer is lacking. In addition, although previous study has suggested that tumor microbial communities are largely unique to each cancer type and microbial profiles can be used to distinguish cancers ^2^, whether both pan-cancer and cancer-type-specific correlations between the tumor microbiome and host molecular features exist are not clear.

Recent advances have shown that the integration of multi-omics data, which include genomics, epigenomics, transcriptomics, and proteomics, can help us better understand how the host and microbe interact by correlating host molecular aberrations with the microbiota ^12–14^. The Cancer Genome Atlas (TCGA), one of the largest cancer sequencing efforts, has profiled host molecular aberrations at the DNA, RNA, protein and epigenetic levels for 32 cancer types. The tumor microbiome in the entire TCGA cohort has also been systematically characterized ^1, 2^. Hence, it is possible and imperative to link the pan-cancer microbiome and host multi-omics data to gain new insights into potential functional roles of the tumor microbiome in the cancer ecosystem.

Here, we comprehensively characterized host-microbe associations in 32 cancer types using multi-omics profiles from TCGA. Our analysis identified both cancer-type-specific and pan-cancer host-microbe associations. We validated and investigated several recurrent host-microbe associations with certain molecular and clinical features. Finally, we present MOMAC (Multi-Omics and Microbiome Associations in Cancer), an online web portal (https://comics.med.sustech.edu.cn/momac) for exploring host-microbe associations identified in pan-cancer.

## Results

### Identifying host-microbe associations at the pan-cancer and multi-omics level

In this study, we integrated tumor microbiome profiles consisting of 1,206 bacteria, 224 fungi, 138 viruses and 62 archaea, along with host molecular features including 10,216 gene expression, 30,716 CpG island methylation, and 193 protein expression, for nearly 10,000 samples across 32 cancer types collected from TCGA. To model the cancer ecosystem, we also integrated proportions of 20 cell types derived from the deconvolution of bulk transcriptomics data. To evaluate the associations between the tumor microbiome and host features, we employed various analytical techniques, including sparse canonical correlation analysis (sparse CCA), Spearman’s rank correlation test and the randomized lasso regression analysis (**Figure 1A**).

**Figure 1.**
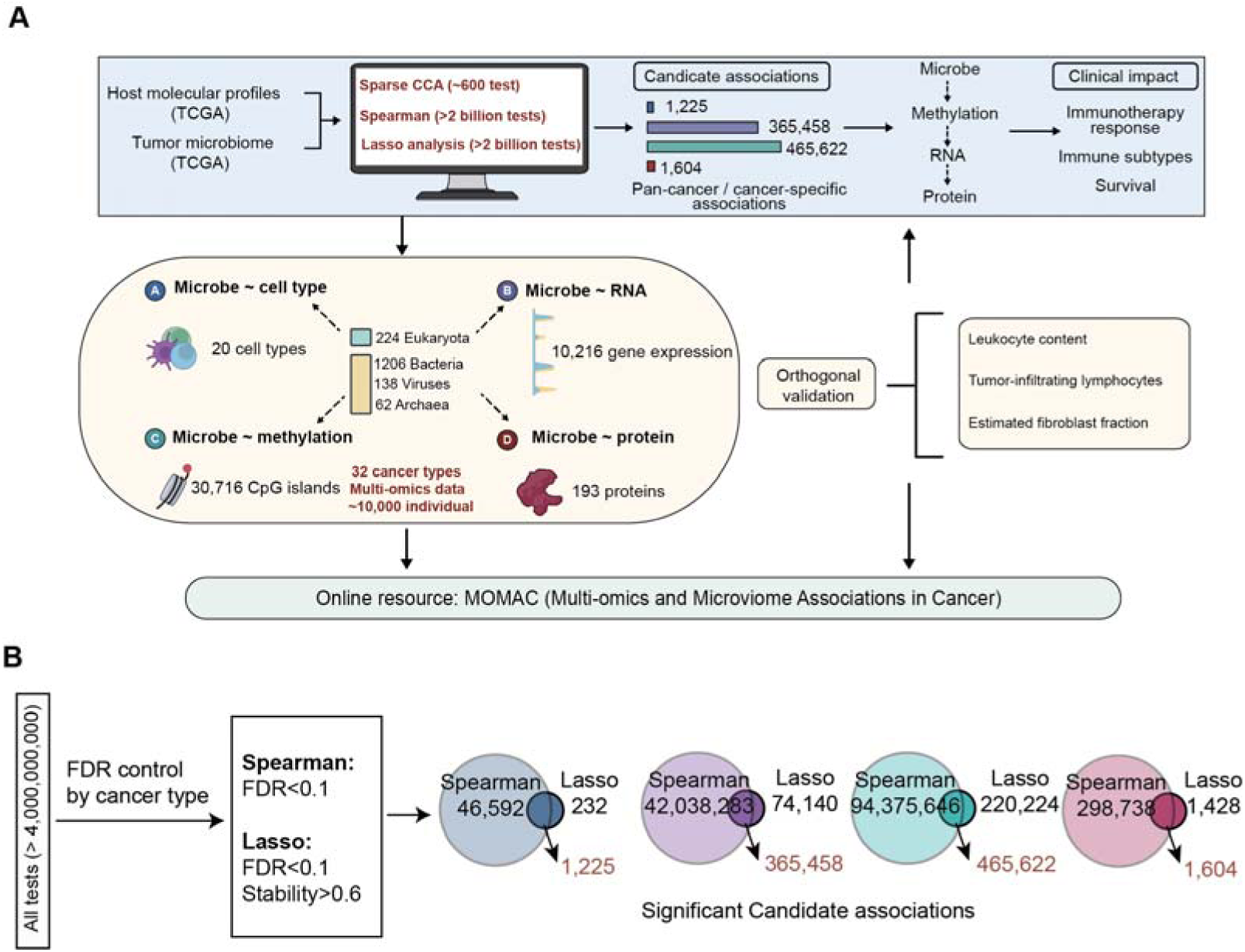
Overview of the study. **A)** Workflow for the identification and characterization of associations between host molecular features and the tumor microbiome at pan-cancer and multi-omics level. **B)** Selection of significant candidate associations for host cell types (blue), gene expression (purple), CpG island methylation (green) and protein expression (red). Final numbers of candidate associations are shown as the overlap of Spearman’s rank correlation test and lasso regression test.

After screening more than 4 billion microbe and host feature pairs, we identified 836,738 candidate associations (Benjamini-Hochberg adjusted p-value < 0.1 for both the Spearman’s test and the lasso test, and stability > 0.6 for the randomized lasso test; **Methods**), which accounted for only 0.0198% of the total association tests. Specifically, we discovered 1,225, 365,458, 465,662, and 1,604 associations between microbes and cell types, gene expression, CpG island methylation, and protein expression, respectively. Our results demonstrated the feasibility of identifying associations between the tumor microbiome and host features using multi-omics data in pan-cancer, and such associations can be found at different omics and cancer types.

### Validating known associations on pathogenic infections

To assess whether our candidate associations captured known host-microbe interactions, we examined microbes that were causally linked to cancer. We first analyzed patients with liver hepatocellular carcinoma (LIHC) who may be infected with the Hepatitis B virus (HBV species). LIHC patients had an overabundance of *Orthohepadnavirus* (HBV genus), which is a major causative etiological factor. Overall, 13 consistent hub genes identified by our work and recent studies ^15–17^ were confirmed to be associated with HBV infection (**Figure S1A, Table S1**). Our gene set overrepresentation analysis using gene ontology (GO) and Kyoto Encyclopedia of Genes and Genomes (KEGG) revealed that genes associated with the HBV genus were enriched in biological pathways like mitotic nuclear division, cell cycle, DNA replication, chromosome segregation and nuclear division, which were also consistent with previously identified pathways that may contribute to the pathogenesis of HBV-infected liver cancer (**Figure S1A**)^17^.

Next, we analyzed *Helicobacter pylori* (within the *Helicobacter* genus) in stomach adenocarcinoma (STAD) patients, which is known to be associated with gastric carcinogenesis through p53 degradation ^18^. Among 13 genes associated with the *Helicobacter* genus, nine genes were also identified in previous studies (**Figure S1B, Table S1**) ^19^. These genes were involved in GO terms related to homophilic cell adhesion via plasma membrane adhesion molecules and epithelial tube morphogenesis, and KEGG pathways related to cancer (**Figure S1B**). We also discovered negative correlation (*r* = −0.195, *p* = 0.0025) between *Helicobacter* and *TP53* gene expression (**Table S1**). Collectively, these results indicate that our candidate associations could capture known associations between host molecules and oncogenic microbes.

### Identifying recurrent associations between tumor microbiome and host features

To provide a more detailed insight into the distribution of candidate associations, we categorized them into cancer-type-specific and pan-cancer associations as well as gastrointestinal tract (GI) cancer and non-GI cancer associations (**Figure 2A**). Pan-cancer associations were defined as recurrent host-microbe associations if they were found in at least eight out of 32 cancer types for cell types, gene expression and CpG island methylation, and at least four cancers for protein expression. Our analysis revealed that approximately 24.98%, 54.84%, 79.93% and 86.16% of associations for cell type, RNA, methylation and protein were found only in one cancer type (cancer-type-specific), while 36.57%, 11.13%, 0.98% and 3.05% associations were recurrent pan-cancer associations, respectively. We also noticed that seven GI cancers (colon adenocarcinoma, COAD; cholangiocarcinoma, CHOL; esophageal carcinoma, ESCA; pancreatic adenocarcinoma, PAAD; rectum adenocarcinoma, READ; STAD; LIHC) accounted for 16.77%-24.47% of all associations for different host feature types. For the rest of the manuscript, we focused on characterizing recurrent pan-cancer associations, while providing examples of cancer-type-specific associations. Notably, all potential host-microbe interactions can be explored via the MOMAC web portal.

**Figure 2.**
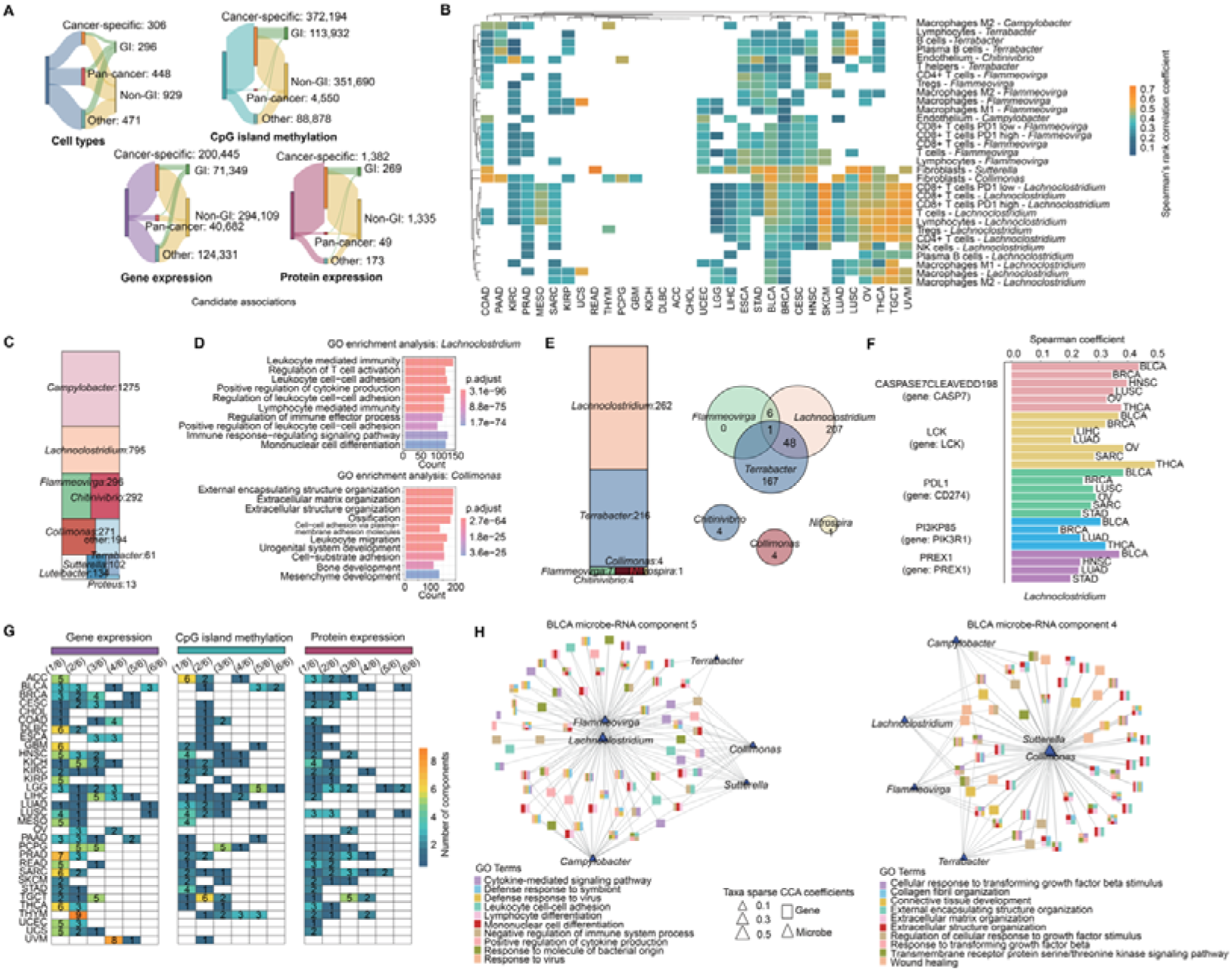
Pan-cancer recurrent host-microbe associations. **A)** The Sankey diagram shows candidate associations grouped by recurrence or organ system. Recurrence grouping includes cancer-specific (only found in one cancer type), pan-cancer (>=4 cancer types for protein and >=8 cancer types for the rest) or other. Organ system grouping includes gastrointestinal (GI) cancer and non-GI cancer. GI cancer includes COAD, CHOL, ESCA, LIHC, PAAD, READ and STAD. **B)** Heatmap shows pan-cancer associations between microbes and cell types. Color shows Spearman’s correlation coefficients. **C)** Mosaic plot shows the top ten microbes with the highest number of associated gene expression. **D)** Gene ontology overrepresentation analysis for two representative microbes, *Lachnoclostridium* and *Collimonas*. **E)** Mosaic plot on the left shows the number of CpG islands associated with pan-cancer microbes and the Venn diagram on the right shows the overlap of associated CpG islands for pan-cancer microbes. **F)** Barplot shows pan-cancer associations between *Lachnoclostridium* and protein expression, with gene names showed in brackets. **G)** Heatmap shows the number of sparse CCA components for six pan-cancer key microbes. Columns 1 to 6 correspond to the number of pan-cancer key microbes co-occurred in the same component. **H)** Network plots show two examples in BLCA for microbe-RNA sparse CCA components. Squared nodes in the network represent genes colored by their GO terms, and triangular nodes are microbes. The size of triangle represents the absolute value of sparse CCA coefficients for microbes. Only microbe-RNA associations with Spearman’s rho > 0.3 and top 10 GO terms based on adjust *p* values are showed.

Interestingly, for host cell type associations, we discovered a large proportion of cancer types (28/32, 87.5%), except for four cancers with limited sample size (**Figure 2B, Figure S2A-B**), exhibited strong associations between microbes and non-tumor cells in the tumor microenvironment (TME). Non-tumor cells such as immune cells and fibroblasts, are crucial components of the TME, affecting tumor immune reaction and response to immunotherapy ^20, 21^. Moreover, we identified two clusters of recurrent pan-cancer associations between the tumor microbiome and non-tumor cells: cluster 1 between *Lachnoclostridium*, *Flammeovirga*, *Terrabacter* and *Campylobacter* and immune cells, and cluster 2 between *Collimonas* and *Sutterella* and fibroblasts (**Figure S2C**). Hereafter, these six microbes were referred to as pan-cancer key microbes.

Microbes within these two clusters were also associated with many other host molecular aberrations. At the gene expression level, for example, 1,275 and 795 genes were discovered to be associated with *Campylobacter* and *Lachnoclostridium*, respectively, which were among the top ten microbes having the highest number of associations with host gene expression (**Figure 2C, Table S2**). We also annotated gene pathways associated with two representative microbes from two clusters, *Lachnoclostridium and Collimonas*. *Lachnoclostridium* was discovered to be correlated with activation and adhesion of immune cells, while *Collimonas* was associated with extracellular organizations (**Figure 2D**). Gene set enrichment analysis for the other four microbes can be found in **Figure S3A**, and the results were consistent with their non-tumor cell associations. Namely, cluster 1 microbes mainly correlated with host immune responses and cluster 2 microbes mainly correlated with extracellular organizations. Similarly, the DNA methylation status of 262 and 199 CpG islands were associated with *Lachnoclostridium* and *Terrabacter*, respectively. We also found that microbes correlated with immune cells in cluster 1 tend to be associated with a shared set of CpG islands (**Figure 2E, Table S3**). By linking CpG islands to genes, we found cluster 1 microbes were associated with pathways like actin organization and cell adhesion (**Figure S3B**). For *Lachnoclostridium*, we also identified 15 genes with both mRNA and methylation associations, and these genes were related to the immune signaling (**Figure S3C**). Interestingly, we also noticed some microbes associated with multiple CpG islands linked to the same gene. For example, *Lachnoclostridium* was found to have multiple associated CpG islands linked to *AGAP2*, *NCOR2*, *HIVEP3*, and *HLA-F*, while four CpG islands on *DOCK1* correlated with *Terrabacter* abundance (**Figure S3D**). Since only 66.9% of TCGA samples having the protein expression data, we detected merely eight recurrent associations, mainly (n = 5) for *Lachnoclostridium* (**Figure 2F, Table S4**).

To further explore the role of pan-cancer key microbes, we employed the sparse CCA method to study the co-occurrence and host association patterns of them. In most cancer types, more than two of these six microbes could be observed in the same component (**Figure 2G, Table S5**). In several instances, we found all six microbes belonging to the same component, including associations between gene expression and microbes in bladder urothelial carcinoma (BLCA), lung adenocarcinoma (LUAD) and lung squamous cell carcinoma (LUSC), between CpG island methylation and microbes in BLCA and brain low grade glioma (LGG), and between protein expression and microbes in BLCA, LGG and LUSC. The sparse CCA analysis also allowed us to compare the relative contribution of different microbes in certain pathways. For example, in the fifth sparse CCA component of BLCA, we highlighted the pivotal role of cluster 1 microbes in comparison with cluster 2 microbes in host immune signaling pathways (**Table S6, Figure 2H**), as the absolute coefficients for *Lachnoclostridium* (−0.56), *Flammeovirga* (−0.26) and *Campylobacter* (−0.24) were higher than *Collimonas* (−0.10) and *Sutterella* (−0.11); in contrast, we noticed a higher contribution of *Collimonas* (−0.48) and *Sutterella* (−0.48) to extracellular organizations than *Lachnoclostridium* (−0.17), *Flammeovirga* (−0.17), *Campylobacter* (−0.17) and *Terrabacter* (−0.16) in the fourth sparse CCA component (**Table S6**). Similar patterns of differential contribution of cluster 1 and cluster 2 microbes in host pathways were also observed at the DNA methylation and protein expression levels using sparse CCA (**Figure S4, Table S7-8**). Altogether, these results demonstrated that recurrent host-microbe associations could be identified in pan-cancer and were mainly related to non-tumor cells and relevant signaling pathways in the TME.

### Validating recurrent associations using orthogonal data

Recurrent host-microbe associations identified above used cell type data based on RNA-Seq, thus we wondered if they can be validated by cell type estimations based on orthogonal data types. For the purpose of validation, we selected two control microbes (*Bacteroides*, a common bacterium, and *Neisseria*, a classified pathogen) that had no associations with cell types in the initial test and whose average abundance was similar to the pan-cancer microbes. Firstly, we used leukocyte fraction estimated from DNA methylation ^22^, and found that cluster 1 microbes but not control microbes showed consistent associations with the leukocyte fraction across multiple cancer types (**Figure 3A**). Next, we examined the association between microbes and tumor-infiltrating lymphocytes based on Hematoxylin and Eosin (H&E) images for 13 tumor types ^23^. As expected, the results were consistent with microbe and cell type associations based on RNA-Seq for *Lachnoclostridium*, *Flammeovirga* and control microbes (**Figure S5**). For example, for *Lachnoclostridium*, in 8 out of 9 tumor types showing associations with immune cells previously, we confirmed again the association between microbe and tumor-infiltrating lymphocytes, whereas this correlation was absent in three out of four cancer types lacking associations in previous analysis (**Figure 3B**). To further validate cluster 2 microbes associated with fibroblast, we defined an estimated fibroblast fraction, approximated by the non-tumor and non-leukocyte fraction of the TME (**Methods**). We again observed expected associations for cluster 2 microbes in cancer types where it had been initially discovered, but not in other cancer types nor in control microbes (**Figure 3C**). Collectively, our validation analysis indicated that recurrent pan-cancer host-microbe associations with immune cells and fibroblasts could be validated by orthogonal data.

**Figure 3.**
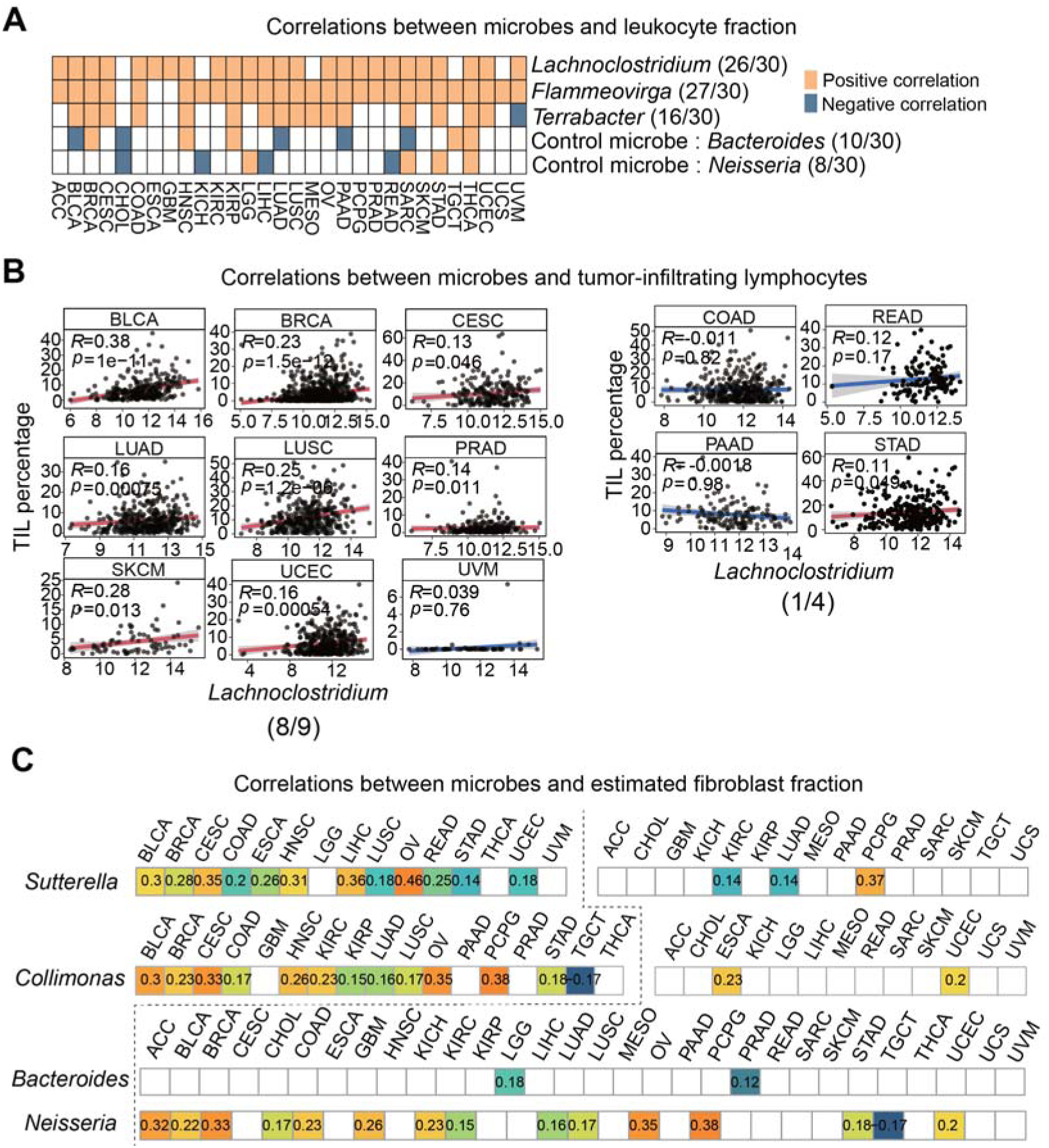
Validation of associations between pan-cancer key microbes and host cell types. **A-B)** Validation of associations between pan-cancer immune-related microbes (cluster 1) and leukocyte fraction from methylation (A), and tumor-infiltrating lymphocytes from pathological images (B). For B, the left shows nine cancer types with significant associations in the initial test, and the right shows four cancer types that were not; the red and blue fitted lines represent significant (*p* < 0.05) and non-significant Spearman’s rank correlations in the validation analysis, respectively. **C)** Validation of associations between fibroblast-related microbes (cluster 2) and estimated fibroblast fraction. Numbers are the Spearman’s rank correlation coefficients. White squares represent no correlation (*p* > 0.05).

### Identifying putative microbe-methylation-RNA-protein axis

After analyzing host-microbe associations at the single omics level, we performed an integrative analysis to identify host-microbe associations across multiple omics, which may suggest key genes and how these genes are regulated by the tumor microbiome. As expected, we observed a high prevalence of overlaps between gene expression and proteins (**Figure 4A**). For instance, 41.67% (n = 53) microbe-protein associations in ovarian cancer (OV) were also detected at the RNA level. Furthermore, the *Lachnoclostridium* genus was found to be correlated with mRNA and protein levels of *LCK*, *CD274* and *PREX1* in multiple cancers. Interestingly, we observed varied levels (0-2,022 genes) of overlaps between microbe-RNA and microbe-methylation associations across 32 cancer types, suggesting the possibility of the tumor microbiome contributing to host gene expression via affecting CpG island methylation in certain cancers (**Figure 4B**).

**Figure 4.**
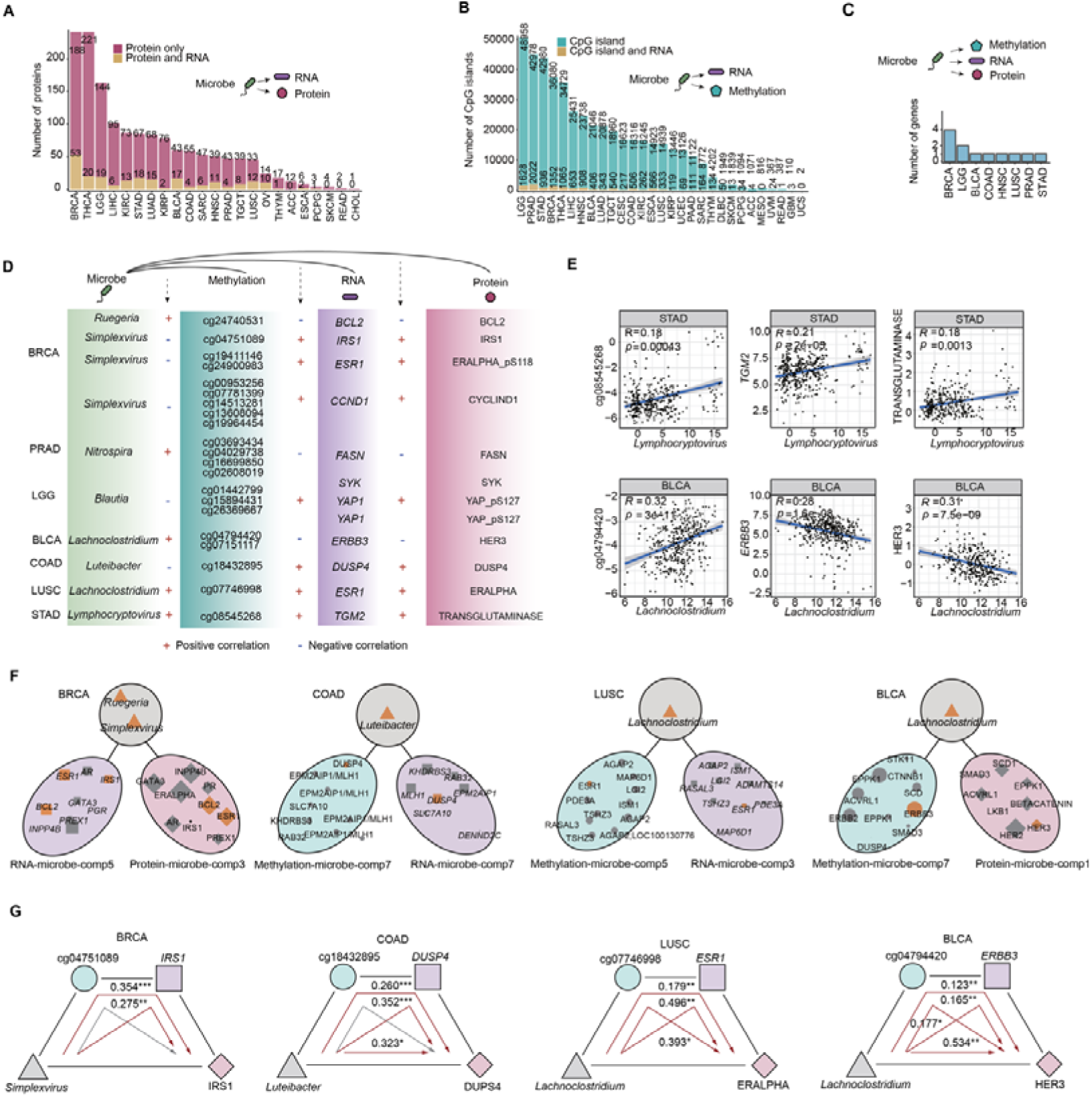
Overlap of multi-omics between specific microbes and host molecular aberrations. **A-C)** Bar plots show the number of overlaps for associations among RNA-protein (A), methylation-RNA (B) and methylation-RNA-protein (C). **D)** Associations between microbes and host genes with complete microbe-methylation-RNA-protein paths. **E)** Scatterplots show two multi-omics association examples from D. The Spearman’s rank correlation coefficients and *p* values are showed. **F)** Microbes associated with the same sets of genes across sparse CCA components (comp) of different omics. CpG islands have been linked to genes. The size of rectangular nodes indicates the absolute value of coefficients from sparse CCA. **G)** Sequential mediation analysis results for tumor microbes and host genes. Each path represents a potential microbe-methylation-RNA-protein axis from D (also in Figure S7). The proportional effect size (shown in number) and *p* values (*, <0.05; **, <0.01; ***, <0.001) are showed in each plot, and the red line represents significant path with *p* values < 0.05 and non-zero proportional effect size.

In addition, we discovered 12 genes with associations across all three omics in eight cancers (**Figure 4C**). Intriguingly, although the direction of associations for microbe-RNA and microbe-protein was always consistent, the direction for microbe-methylation associations was not (**Figure 4D**), which may be related to the position of CpG islands (gene body or promoter) and the context-dependent manner of gene regulation by promoter hypermethylation ^24^. For example, *Lymphocryptovirus* (Epstein–Barr virus) was enriched in EBV-infected primary STAD (**Figure S6A**), and positively correlated with T cells especially CD8+ T cells in our study (**Figure S6B**), which were consistent with previous studies^25, 26^. Our results suggested that *Lymphocryptovirus* was associated with promoter hypermethylation (cg08545268, r = 0.18, *p* = 0.00043) and increased mRNA (*r* = 0.21, *p* = 2×10^−5^) and protein (*r* = 0.18, *p* = 0.0013) levels of *TGM2*, a known marker gene for worse prognosis in stomach cancer ^25^. Additionally, the overexpression of *ERBB3* and its protein (HER3) have been reported in BLCA, which was involved in tumor progression and metastasis ^27^. Our results implied that *Lachnoclostridium* might inhibit *ERBB3* expression (RNA: *r* = −0.28, *p* = 1.6×10^−8^, protein: *r* = −0.31, *p* = 7.5×10^−9^).

Next, we employed sparse CCA, a complimentary method with the advantage to identify cross-omics association modules. We observed that several microbe-RNA modules were also captured by microbe-protein and/or microbe-methylation analysis. For instance, in breast invasive carcinoma (BRCA), microbes including *Ruegeria* (coefficient = −0.33), *Simplevirus* (coefficient = −0.41), and genes including *ESR1* (coefficient = 0.07), *IRS1* (coefficient = 0.04) and *BCL2* (coefficient = 0.08) were members of the fifth sparse CCA component for microbe-RNA associations, and the same set of microbes and genes also contributed (absolute coefficient = 0.02-0.09) to the third component of microbe-protein associations (**Figure 4F, Table S9**). In COAD, LUSC and BLCA, we also found two microbes (*Luteibacter* and *Lachnoclostridium*) that may affect the same features in different omics (**Figure 4F, Table S9**).

Integrative analysis above further supported the idea that the tumor microbiome may modulate host molecular features via the microbe-methylation-RNA-protein axis, so we next performed the sequential mediation analysis, a statistical method that can test if one variable (microbe) affects another (RNA/protein) via a mediator (methylation). In BRCA, we connected the *Simplexvirus* genus of herpes simplex virus (HSV) with methylation, RNA, and protein changes for *IRS1* and *ESR1* (**Figure 4G, Figure S7** and **Table S10**). We speculated that *Simplexvirus* might enhance both the gene and protein expression of *IRS1* and *ESR1* via hypomethylating two corresponding CpG islands marked by cg04751089 and cg24900983, respectively. Moreover, *Blautia* and *Luteibacter* might contribute to the hypomethylation of *SYK* (promoter), *YAP1* (promoter), and *DUSP4* (gene body) to modulate their expression in LGG (**Figure S7**). In prostate adenocarcinoma (PRAD) and BRCA, similar microbe-methylation-RNA-protein axis was found for *Nitrospira*-*FASN* and *Ruegeria-BCL2* (**Figure S7**). For pan-cancer key microbes, *Lachnoclostridium* was associated with the hypermethylation of cg07746998 (*ESR1*) and cg04794420 (*ERBB3*), and their corresponding gene/protein levels in LUSC and BLCA, respectively (**Figure 4G**). Collectively, through multi-omics integrative analysis, our results indicated that microbes could potentially affect host gene and/or protein expression via altering genome methylation.

### Investigating the clinical impact of microbes with pan-cancer associations

We next sought to characterize the clinical impact of these microbes having recurrent pan-cancer associations. Since the gut microbiota can shape the response to immune checkpoint inhibitors ^28, 29^, we suspected that the tumor microbiome may also be associated with immunotherapy response. One previous study reported that the T cell-inflamed gene expression profile (GEP) score can be utilized to classify responders versus non-responders to PD-1 checkpoint blockades ^30^. Hence, we calculated GEP scores for all samples and tested their associations with the tumor microbiome. Our analysis revealed significant positive associations between the GEP score and six pan-cancer microbes across multiple cancer types (**Figure 5A**), especially for *Lachnoclostridium* (*r* = 0.31-0.70), *Flammeovirga* (*r* = 0.30-0.54), and *Terrabacter* (*r* = 0.23-0.42).

**Figure 5.**
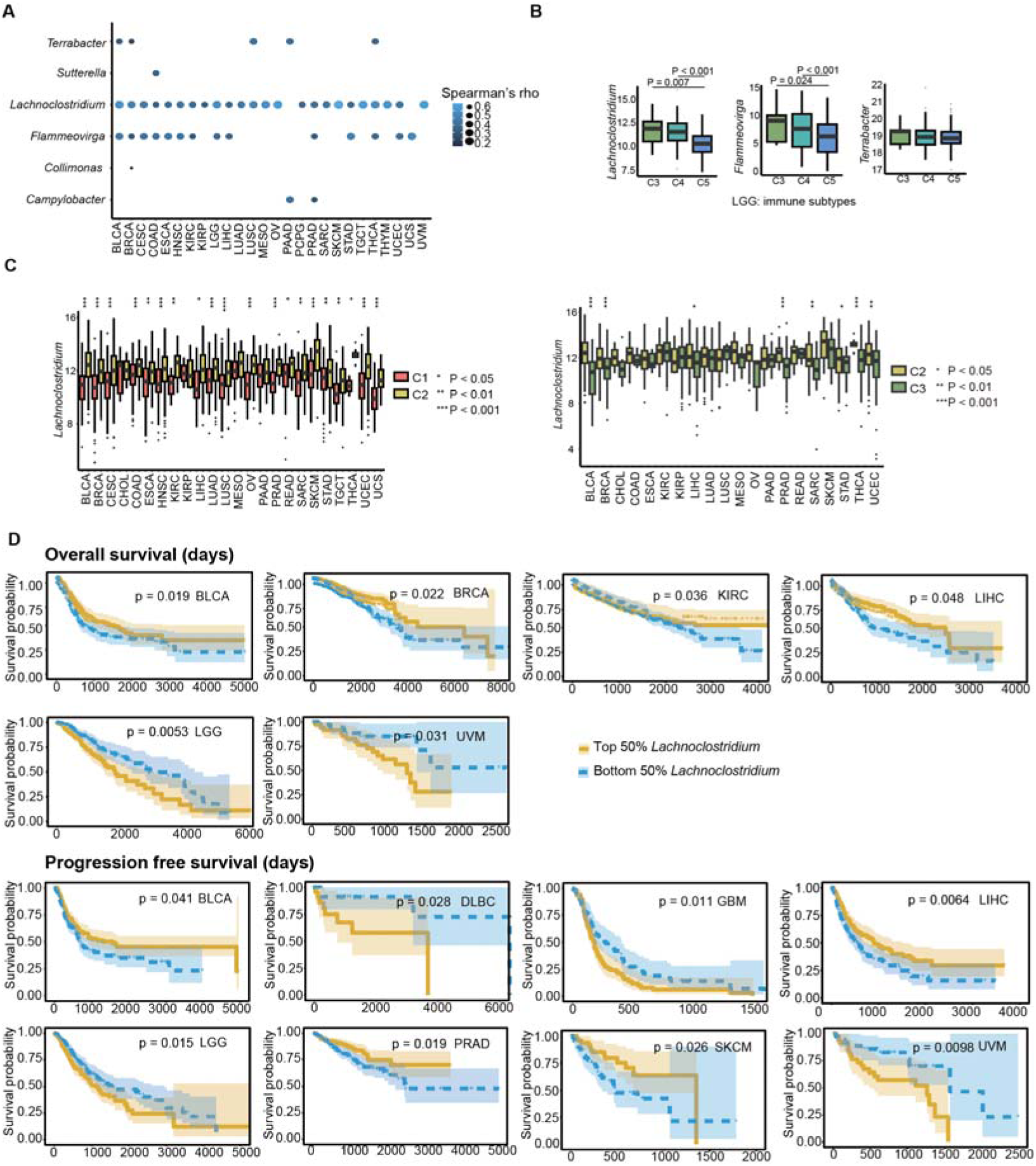
Clinical impact of pan-cancer key microbes. **A)** Associations between microbes and GEP scores. Significant associations are shown with circles (Spearman: FDR < 0.1; lasso: FDR < 0.1, and stability selection > 0.6). **B-C)** Associations between *Lachnoclostridium* and immune subtypes in LGG (B) and pan-cancer (C). C1, wound healing; C2, IFN-γ dominant; C3, inflammatory; C4, lymphocyte depleted; C5, immunologically quiet. Wilcoxon tests were used to determine statistical significance (* *p* < 0.05, ** *p* < 0.01, *** *p* < 0.001). **D)** Kaplan-Meier plots show the potential clinical impact of *Lachnoclostridium* in OS and PFS across multiple cancer types. *P* < 0.05 was considered as significant.

All TCGA tumors have been divided into six immune subtypes (C1 to C6) with clinical relevance in a previous study ^22^, and we next examined if our pan-cancer key microbes contributed to tumor immune subtypes. The C5 (immunologically quiet) subtype featured by few lymphocytes is mainly reported in LGG ^22^; thus, we first compared the abundance of immune-related microbes between C3 (inflammatory), C4 (lymphocyte depleted) and C5 in LGG (**Figure 5B**). As expected, the abundance of *Lachnoclostridium* and *Flammeovirga* significantly reduced in C5 compared with C3 (*p* = 0.007 and 0.024, respectively) and C4 (both *p* < 0.001) in LGG. Moreover, C1 (wound healing), C2 (IFN-γ dominant) and C3 (inflammatory) are present in most cancers ^22^, and we found significant differences in the abundance of *Lachnoclostridium* between these immune subtypes in multiple cancer types (**Figure 5C**). Note that *Lachnoclostridium* was also positively correlated with the *IFNG* expression (C2 immune subtype marker) in 18 cancer types (**Table S12**).

Finally, these findings with immunotherapy response and immune subtype prompted us to investigate the association between pan-cancer key microbes and prognosis. We defined top and bottom groups based on the median abundance of pan-cancer key microbes and compared the overall survival (OS) and progression-free survival (PFS) between these two groups. Stratification of patients based on the abundance of *Lachnoclostridium* was significantly associated with OS and PFS in multiple cancer types (**Figure 5D**), including positive correlations with OS in BLCA, BRCA, kidney renal clear cell carcinoma (KIRC) and LIHC, and PFS in BLCA, LIHC, PRAD and skin cutaneous melanoma (SKCM), and negative correlations in LGG (OS: *p* = 0.015, PFS: *p* = 0.02) and GBM (glioblastoma multiforme; PFS: *p* = 0.0098), UVM (uveal melanoma; OS: *p* = 0.021, PFS: *p* = 0.0098) and DLBC (diffuse Large B-cell Lymphoma; PFS: *p* = 0.028). We also identified significant associations with OS and/or PFS for other pan-cancer key microbes (**Figure S8**). Together, these results suggested that the tumor microbiome has potential clinical utility such as patient stratification and prediction of therapeutic response and prognosis.

## Discussion

In this study, we linked the tumor microbiome and host molecular features at the pan-cancer and multi-omics level. We screened the correlations and further characterized six microbes with recurrent pan-cancer associations in detail. Note that there were also numerous cancer-type-specific host-microbiome associations that could be explored via our MOMAC web portal. Additionally, we have undertaken extensive effort to validate our findings and correlate them with clinical features. The prevalence and importance of the tumor microbiome has been indicated in multiple previous studies ^31–34^; our systematic study here can improve our understanding of the host-microbe interplay in cancer and suggest potential molecular mechanisms.

Our observations confirmed several known host-microbe associations. The HBV genus is causally related to LIHC^35^. Previous studies have discovered that HBV could integrate at the *TERT* and *MLL4* genes, increasing their expression, which contributes to carcinogenesis ^35–37^. In our study, we found that the *TERT* expression and the *MLL4* hypomethylation was positively and negatively associated with the HBV abundance, respectively. However, some cancer-related genes such as *PTPRD*, *UNC5D*, *NRG3*, *CTNND2* and *AHRR* that are altered in HBV-infected liver cancer were not identified in our study, suggesting future studies are required to collect more samples and/or improve the method for host-microbe association study. In STAD, *Helicobacter pylori* infection is known to trigger a proinflammatory response and enhance p53 degradation in host cells due to the presence of CagA protein ^38, 39^, which is also consistent with our analysis. Moreover, previous study reports that the promoter hypermethylation of *MGMT* is associated with *Helicobacter pylori* CagA-positive strains ^40^, and we also find positive correlation between overabundant *Helicobacter* and the hypermethylation of *MGMT*. Thus, our analysis recaptures known host-microbe interplays and further suggest the possibility of finding new host-microbe associations through MOMAC.

From our pan-cancer findings, we speculate that the tumor microbiome may contribute to tumor progression and development via affecting the host immune responses and cancer-related signaling pathways. A recent study reports that tumor microbes belonging to the Lachnospiraceae family can enhance the activation of CD8+ T cells ^13^. Similarly, in COAD, we discovered positive association between *Lachnoclostridim* (within the Lachnospiraceae family) and T lymphocytes. Besides *Lachnoclostridium*, we discovered other microbes associated with immune cells in multiple cancers, including *Flammeovirga*, *Terrabacter* and *Campylobacter*. Of note, the *Campylobacter* has also been reported recently in PDAC to be related to immune cells ^41^. Furthermore, we also identified two microbes associated with fibroblasts and pathways related to extracellular matrix. Hence, non-tumor cells in the TME might be important targets for pan-cancer microbes and future studies should be conducted to validate and study such interplays *in vivo* and *in vitro*.

Furthermore, our multi-omics integration using multiple algorithms shows strong evidence of associations between tumor microbes and host molecular profiles. The CpG island hypermethylation and hypomethylation are common epigenetic signals that regulate gene expression ^42^. Previous studies have shown that the gut microbiome has an impact on host DNA methylation, which in turn contributes to colon cancer development ^43^. Here we also tested the probability of the tumor microbiome modulating host gene expression via CpG island methylation by sequential mediation analysis. Our results indicate that the microbe-methylation-RNA-protein axis can be observed for multiple cancer-related genes across a wide range of cancer types, although how tumor microbes induce host DNA methylation changes requires more studies. Previous studies demonstrated that butyrate produced by gut microbes can alter DNA methylation in cell lines ^44^, so if certain toxins produced by tumor microbes have similar functions should be tested experimentally in the future.

In conclusion, we conduct the first pan-cancer multi-omics association study between host molecular features and the tumor microbiome, and identify potential associations with putative molecular mechanisms within the TME which can be further explored in our MOMAC web portal.

### Limitations of the study

In this study, we discovered candidate associations using the TCGA cohort, one of the most comprehensive tumor sequencing efforts, while additional large multi-omics cohorts should nonetheless be used in future studies to validate our findings and explore potential environmental and/or genetic factors that may shape these associations. We mainly focused on investigating microbes with recurrent pan-cancer associations here. Nevertheless, there are more cancer-type-specific associations that are intriguing and may be crucial for cancer biology awaiting characterization. Moreover, we associated the tumor microbiome to host cell types inferred from bulk RNA-Seq, while future work should consider the usage of additional single-cell RNA sequencing and spatially resolved transcriptomics data to directly link the associations between the tumor microbiome and host cells in the cancer ecosystem. Finally, while suggesting potential molecular mechanism, our analysis cannot establish causal links, which demands future functional studies to provide experimental evidence of how the tumor microbiome can alter host DNA methylation, gene and/or protein expression.

## Methods

### TCGA data integration and processing

We closely examined tumor microbiome profiles of 32 cancer types collected from Dohlman et al ^1^ (n = 10,245) and Poore et al. ^2^ (n = 9,751), and batch-corrected data and “Kraken-TCGA-Voom-SNM-All-Putative-Contaminants-Removed-Data” for the tumor microbiome were downloaded from them, respectively. The resulting two microbial tables include 224 fungi, 1,206 bacteria, 138 viruses and 62 archaea, all of which were decontaminated. For host molecular features, RNA (RESM_tpm, n = 10,535), RPPA (n = 7,744), DNA methylation 450K only (n = 9,639) can be downloaded from the UCSC Xena-TCGA Pan-Cancer platform ^45^. Gene names, CpG islands and proteins were paired through the TCGA portal and the official manifest file for Illumina Infinium HumanMethylation450 v1.2 BeadChip. We selected primary tumor and RNA-sequenced samples for further data analysis, resulting in a sample size of 9,600 and 7,936 for Poore et al. and Dohlman et al. datasets, respectively. Note, to ensure data integrity, duplicated samples and multiple samples from the same individual were kept, with only one chosen at random for association testing. Sample sizes for all 32 cancer types can be found in **Figure S2A**. Note that adrenocortical carcinoma (ACC) and sarcoma (SARC) did not have any eukaryotic microbe profiles.

For host gene expression data, we utilized the R package biomaRt to retain only protein-coding genes ^46^. Then, for sparse CCA analysis, to reduce the number of features, we kept only genes expressed in >95% of samples and having variances within the top 75% of all genes by cancer type. For each cancer type’s host methylation, protein, and microbial abundance data, we likewise retained features with variance among the top 75% of all features for the sparse CCA analysis.

### Cell type data collection and processing

First, for initial discovery, we collected bulk RNA-Seq deconvolution profiles for 10,490 TCGA tumors using Kassandra, including abundance estimation of 20 different cell types ^47^. We then took an extensive search of resources to validate our findings, including leukocyte fraction data for 9,783 samples in 30 non-hematologic cancer types based on deconvolution of DNA methylation markers ^22^, and the tumor-infiltrating lymphocyte fraction data for 4,210 samples in 13 cancer types based on AI processing of pathological images ^23^. Since the fibroblast fraction has not been directly evaluated by other methods, we further collected tumor purity data for 10,787 samples in 30 non-hematologic cancer types. After intersecting with the leukocyte fraction and the cancer microbiome data, we calculated an estimated fibroblast fraction for 9,071 samples, which should be positively correlated with the fibroblast fraction in the TME as:

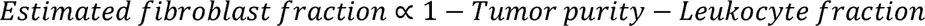

When validating key microbe-cell associations, we included *Bacteroides* and *Neisseria* as common bacteria and pathogen controls, respectively, as they were not associated with cell types in the initial test and the average abundance was similar to that of pan-cancer immune-related microbes.

### Sparse canonical correlation analysis

*Algorithm description*. The sparse CCA algorithm is a multivariate analysis method designed to identify the maximum correlation between variables by linearly projecting two sets of data onto a shared latent space, with regularization to achieve sparsity ^48^. The sparse CCA method has been used in identifying host-microbe associations in literature before ^49^. Given two data matrices with the same number of samples but different number of features, **X***_n_*_x*p*_ and **Y***_n_*_x*q*_, where denoted the number of observations (e.g, tumor samples), *p* and *q* were the number of features (e.g., gene and protein expression, or microbial abundance, etc.), sparse CCA involved finding and *v*, which were loading vectors, that maximize *cor*(*Xu*, *<u>Yv</u>*) In mathematical terms, this optimization problem can be formulated as:

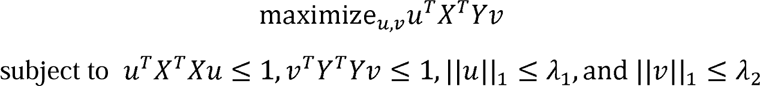

where parameters λ_1_ and λ_2_ correspond to the *L*^1^ norm penalty terms that control the relative weight of the regularization term in the loss function and impacted the solution of the sparse CCA model. These parameters can be adjusted as needed.

#### Grid search for hyperparameters

The range of penalty parameters was constrained to be between 0.1-0.4 using the PMA R package (v1.2.1) ^48^, where a smaller value represents fewer features and a higher level of sparsity. We employed the leave-one-out cross-validation (LOOCV) method to perform a grid search on these hyperparameters (λ_1_, λ_2_), with the objective of determining hyperparameter settings that would yield the highest Pearson’s correlation coefficient between the first pair of canonical variate scores derived from **X***_n_*_x*p*_ and **Y***_n_*_x*q*_ datasets. The correlation was a metric for gauging the efficacy of the chosen hyperparameters in capturing the underlying relationship between the two datasets, ultimately guiding the selection of optimal hyperparameters that maximize the correlation among the canonical variates.

#### Identification of correlated components

We used sparse CCA to obtain highly correlated CCA components for different **X***_n_*_x*p*_-**Y***_n_*_x*q*_ pairs, including microbe-RNA, microbe-methylation, and microbe-protein for each cancer type. Note that, the sparse CCA between fungi and host features were ran separately since the quantification and normalization of fungus abundance were different than other microbes. We utilized optimized hyperparameters to fit a sparse CCA model on each pair of data, and computed the first 10 mutually uncorrelated latent feature spaces (referred to as components ^50^), each encompassing highly correlated data from **X***_n_*_x*p*_ and **Y***_n_*_x*q*_. Then, we evaluated the statistical significance of all components using the LOOCV method, and used the BH procedure to correct for multiple hypothesis testing, defining significant components as those with FDR < 0.1. We employed the R package foreach to construct a parallel computing framework for sparse CCA.

### Spearman’s rank correlation and lasso regression analysis

After filtering out low variable host molecular characters using the methods described above, we used a custom script to calculate the Spearman’s rank correlation between tumor microbes and host molecular features in R. The total number of Spearman’s rank correlation tests is as follows:

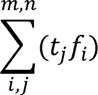

where *f_i_* is the number of features for every host omics phenotype (m = 4, cell types, gene expression, CpG islands and proteins), and *t_j_* is the number of microbes in each cancer type (n = 1,406 for ACC and SARC; n = 1,630 for other cancer types). The resulting *p* values were adjusted using the BH method by cancer and host feature type.

Furthermore, we conducted regression analyses to investigate the association between host molecular features and tumor microbe abundances, using the latter as predictors and the former as outcomes, as described in a recent study ^49^. To mitigate the risk of overfitting, we applied lasso penalized regression model, implemented using the R package glmnet version 4.1.6 ^51^. The lasso regression employs regularization to minimize the following equation:

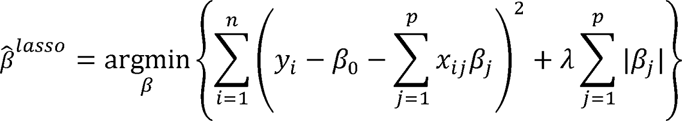

where *n* is the number of samples, *p* is the number of microbes, *y* is host molecular profiles, *x* is tumor microbes and λ is the regularization parameter. To determine the optimal λ value, we utilized LOOCV. Then, *p* values were calculated for the coefficient of each microbe (*β_j_*) using the R package hdi version 0.1-9. We next used the BH method for multiple test correction by cancer and host feature type.

Further, to control for false discoveries, we implemented randomized lasso for stability selection. This approach ran lasso regression 50 times with randomly selected subsets of samples and calculated the fraction of times a feature being selected. A cutoff of 0.6 was used to select stable host-microbe associations. The stability selection was carried out using the R package stabs version 0.6-4 ^52^.

Finally, we screened all associations and created a set of candidate associations. These associations were considered significant if their BH FDR was less than 0.1 for both the Spearman’s rank correlation test and lasso regression test, and if their stability selection value was greater than 0.6. Note that, since host molecular profiles from UCSC Xena have been normalized across pan-cancer, occasionally, some host features in a given cancer type might have constant values (i.e., *y* has no variance), which were excluded in the association analysis. In addition, association analysis was not performed when the usable number of samples was less than 10 in our study.

### Sequential mediation analysis

To integrate multi-omics data and uncover potential biological association pathways, we used R packages bruceR (0.8.10) and lavaan (0.6-15) ^53^ to perform sequential mediation analysis as described in a recent study ^54^. Sequential mediation analysis leverages linear models and bootstrapping techniques to assess if independent variable X (e.g, tumor microbiome) influences dependent variable Y via a seris of mediator variables M_1_ … M_n_, such as host methylation (HostM), RNA expression (HostR) and/or protein expression (HostP). We first conducted the mediation analysis to test the microbe-methylation-RNA axis, by fitting two linear models ^55^:

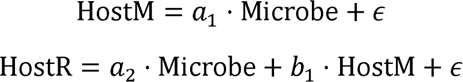

Upon fitting the linear models, the parameters *a*_1_ and *a*_2_ were obtained, and their product (*a*_1_·*a*_2_) represents the average indirect effect of Microbe on HostR. Concurrently, *a*_2_ indicates the average direct influence of Microbe on HostR, adjusting for the average indirect effect mediated by Microbe through HostM.

Likewise, for mediation analysis that includes multiple mediator variables, such as the microbe-methylation-RNA-protein sequential mediation analysis model, the tumor microbiome impacts host methylation (HostM), subsequently affecting host gene expression (HostR), and ultimately influencing host protein expression (HostP). The significance and proportional effect sizes of the mediation effects were determined by fitting the following three linear equations:

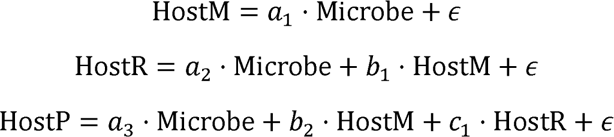

The product of (*a*_1·_*b*_1·_*c*_1_) signifies the average indirect effect of Microbe on HostP mediated by HostM and HostR. The parameter *a*_3_ represents the average direct effect of Microbe on HostP, adjusting for the effects via HostM and HostR.

### Gene set and pathway analysis

We identified gene sets and pathways enriched in genes associated with pan-cancer key microbes and pathogen bacteria. We linked CpG islands to genes using the Illumina official manifest file. We transformed gene SYMBOL to ENTREZID using R packages clusterProfiler and org.Hs.eg.db ^56^, and performed GO (gene ontology) and KEGG (Kyoto Encyclopedia of Genes and Genomes) pathway enrichment analysis with BH correction (pvalueCutoff = 0.25). To visualize host-microbe association pathways, we employed R packages ggraph and scatterpie to construct network graphs between host features and microbes. We also used barplot (showCategory = 10) and cnetplot (showCategory = 5 or 10) in the clusterProfiler package to show the enrichment results.

### GEP score analysis

The T cell-inflamed gene expression profile (GEP) score was calculated based on the sum of 18 inflammatory genes, including *CCL5*, *CD27*, *CD274* (PD-L1), *CD276* (B7-H3), *CD8A*, *CMKLR1*, *CXCL9*, *CXCR6*, *HLA-DQA1*, *HLA-DRB1*, *HLA-E*, *IDO1*, *LAG3*, *NKG7*, *PDCD1LG2* (PDL2), *PSMB10*, *STAT1*, and *TIGIT* as described in literature before ^30^. Further, we tested the associations between GEP scores and microbes based on Spearman’s rank correlation and lasso regression tests. Then, we filtered out significant associations (Spearman: FDR < 0.1; lasso: FDR < 0.1, and stability selection > 0.6).

### Immune subtype analysis

Six immune subtypes spanning multiple tumor types have been classified for 9,127 tumors in TCGA as described in literature ^22^. After overlapping with tumors having microbiome profiles, 9,033 samples were included in the analysis. We compared the abundance of lymphocytes-related microbes between different immune subtypes using the Wilcoxon rank sum test.

### Survival analysis

We obtained curated survival phenotypic data including overall survival (OS) and progression-free survival (PFS) from the UCSC Xena server. We excluded samples with missing OS and PFS data (10,702 samples were retained) ^57^. After paired with the tumor microbiome data, 9,483 (Poore et al. dataset) and 7,800 (Dohlman et al. dataset) samples in 32 cancer types were kept for further analysis. Then, samples were divided into high (top 50%) and low (bottom 50%) groups based on the median abundance of microbes in each cancer type. For MOMAC, the prognostic impact of all microbes on OS and PFS were estimated using Kaplan-Meier analysis and hazard ratios were computed using the R package survminer version0.4.9. In this study, we showed the impact on OS and PFS for six pan-cancer key microbes and *p* values < 0.05 were treated as significant.

### The MOMAC web portal

The web portal can be accessed at https://comics.med.sustech.edu.cn/momac. The platform is composed of five main parts: MOMAC Search, Advanced Search, Survival, Statistics and Download. Significant candidate associations can be queried in “MOMAC Search” by inputting gene, microbe and/or cancer type names. The “Advanced Search” function can be used to select candidate associations with more filter parameters. The “Statistics” page shows basic statistics of candidate associations and the “Survival” page provides survival analysis for a given microbe as described above. Furthermore, all summary statistics of our association analysis can be acquired in the “Download” page.

### Data and code availability

The summary statistics of associations tests have been uploaded to the open-access MOMAC web portal (https://comics.med.sustech.edu.cn/momac), which are available for download to all users. The publicly available microbiome data can be found for fungi (https://github.com/knightlab-analyses/mycobiome/tree/master/Final_files) and other microbes (ftp://ftp.microbio.me/pub/cancer_microbiome_analysis). All code and supplementary data are available via GitHub at https://github.com/comics-bio/pan-cancer-host-microbiome-associations. The corresponding author (Shimin Shuai) can be contacted for further additional information.

## Ethics approval and consent to participate

Not applicable.

## Acknowledgements

C.M., C.S., J.L. and S.S. were partially supported by the National Natural Science Foundation of China (82270239) and the Science and Technology Program of Shenzhen (JCYJ20220530115207016). C.S. and J.Q. were supported by the Guangdong Province Science and Technology Project (2020B1111170014), the Guangdong Basic and Applied Basic Research Foundation (2020A1515010586), the Science and Technology Program of Shenzhen (JCYJ20190809144005609 and JCYJ20210324131212034), the Third People’s Hospital of Shenzhen (G2021014), and the Shenzhen Key Laboratory of Biochip (ZDSYS201504301534057). Computational analysis of this work was supported by the Center for Computational Science and Engineering at Southern University of Science and Technology.

## Contributions

C.M. and S.S. conceived the project; C.M., C.S. and S.S. analyzed the data, interpreted the results, and wrote the manuscript; J.L. designed the web portal; J.Q. and S.S. supervised the project. All authors read and approved the manuscript.

