## Supplementary Information for "Pan-cancer analysis reveals tumor microbiome associations with host molecular aberrations"

**Table S1.** Host molecular features associated with the *Orthohepadnavirus* and *Helicobacter* genus.

**Table S2.** Recurrent pan-cancer associations between RNA and tumor microbes.

**Table S3.** Recurrent pan-cancer associations between CpG island methylation and tumor microbes.

**Table S4.** Recurrent pan-cancer associations between protein expression and tumor microbes.

**Table S5.** The number of six pan-cancer key microbes co-occurred in the same sparse CCA component for different omics and cancer types.

**Table S6.** The coefficients and GO terms of genes and microbes in the fourth and fifth sparse CCA components for BLCA.

**Table S7.** The coefficients and GO terms of CpG islands and microbes in the eighth and tenth sparse CCA components for BLCA.

**Table S8.** The coefficients of proteins and microbes in the eighth and tenth sparse CCA components for HNSC.

**Table S9.** Sparse CCA results for microbes associated with the same sets of genes across components from different omics.

**Table S10.** Sequential mediation analysis for the microbe-methylation-RNA-protein axis.

**Table S11.** Candidate associations between GEP scores and six pan-cancer key microbes in multiple cancers.

**Table S12**. Significant associations between *Lachnoclostridium* and *IFNG* gene expression in multiple cancers.

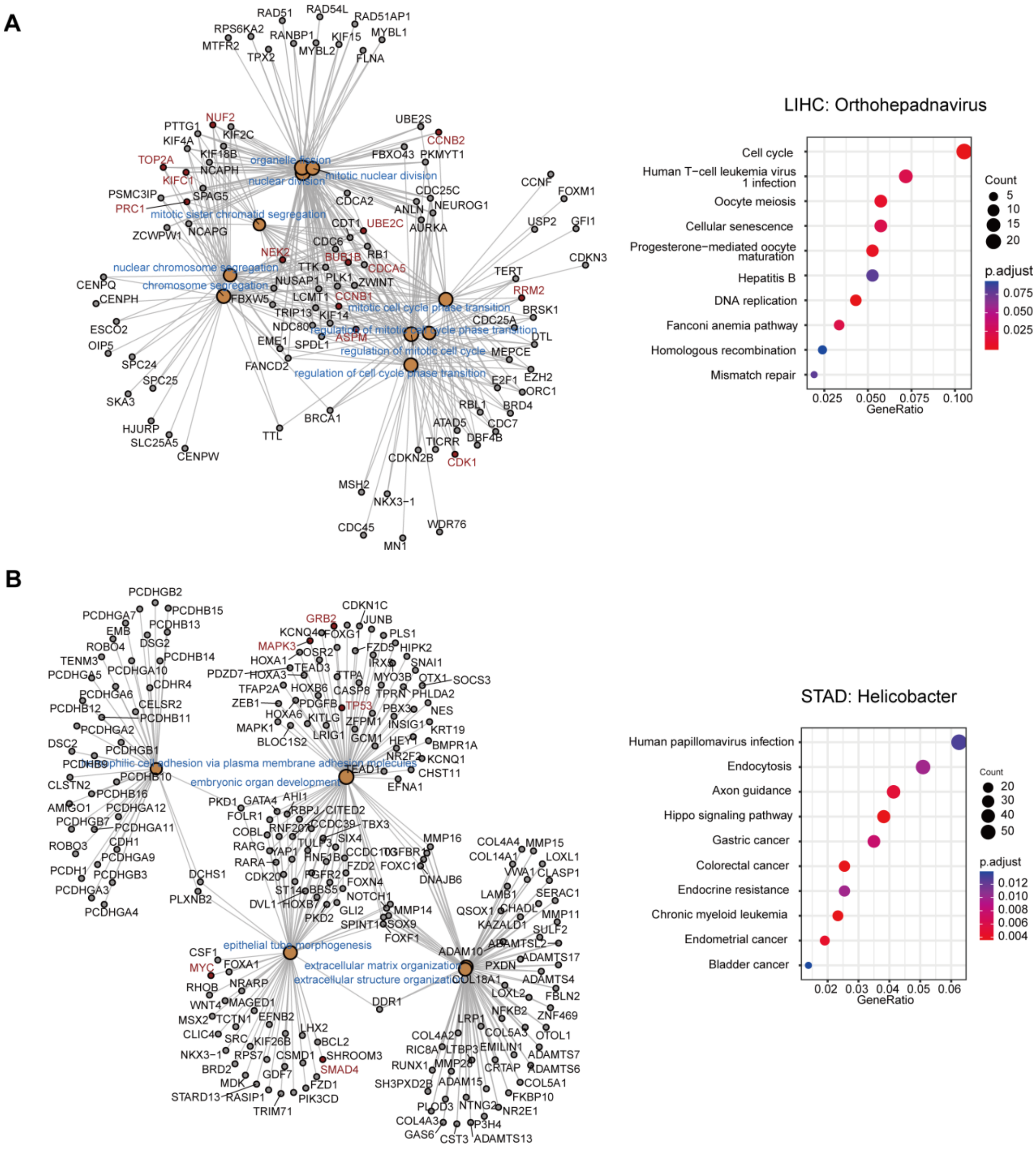


**Figure S1. The GO and KEGG enrichment analysis on confirmed pathogenic infected cancers. A)** ***Orthohepadnavirus*****. B)** ***Helicobacter***. Left panel represented GO enrichment analysis. Panel A showed 10 pathways, and panel B showed five pathways based on adjusted *p* value, and right panel was KEGG enrichment analysis (10 pathways were showed based on gene ratio). Genes colored with red in GO enrichment analysis have been reported and highlighted in previous studies and were discussed in our study.

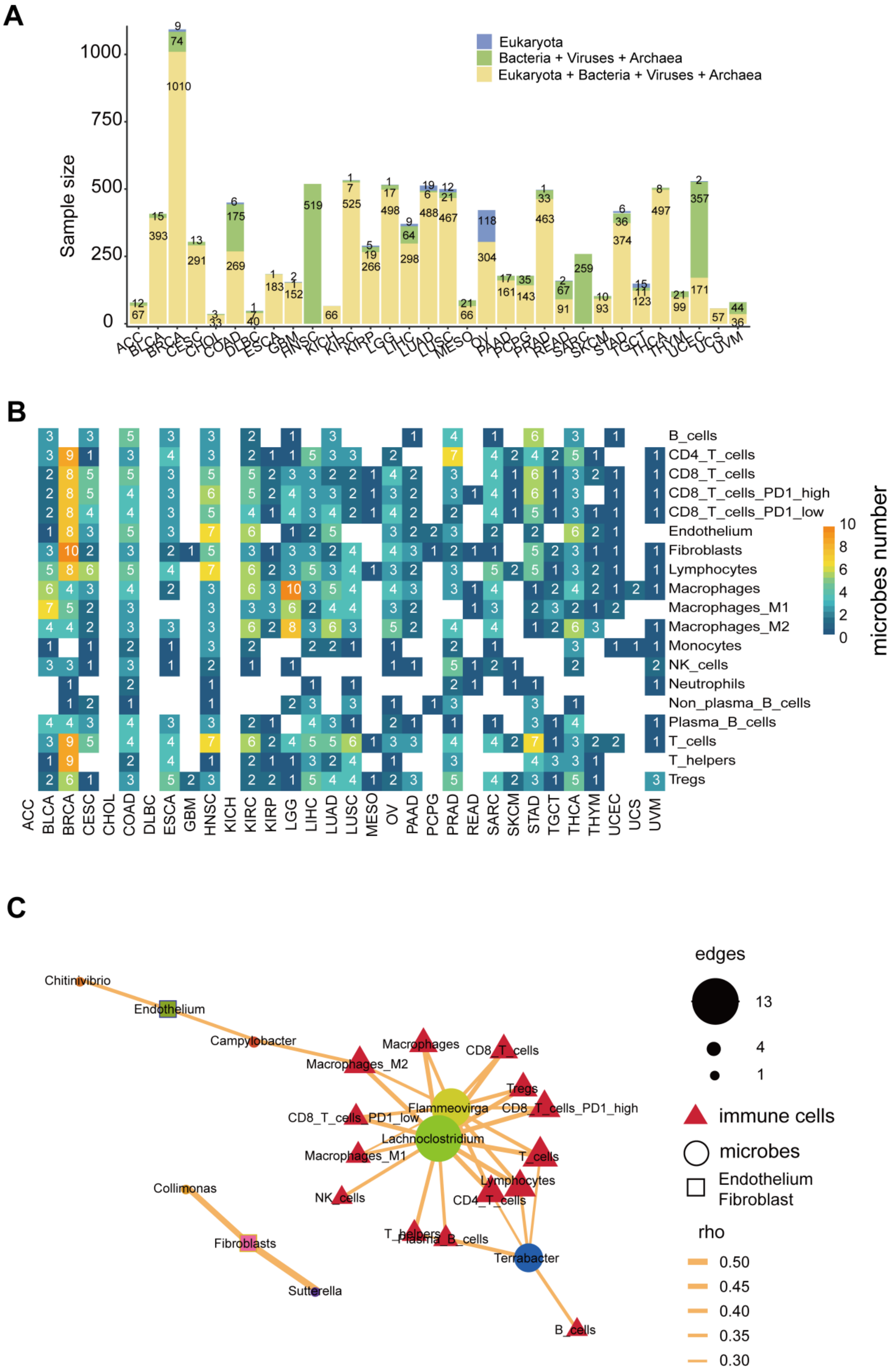
A
Sample size
Eukaryota
Bacteria + Viruses + Archaea
Eukaryota + Bacteria + Viruses + Archaea
ACC BLCA BRCA CESC CHOL COAD DLBC ESCA GBM HNSC KICH KIRC KIRP LGG LIHC LUAD LUSC MESO OV PAAD PCPG PRAD READ SARC SKCM STAD TGCT THCA THYM UCEC UCS UVM
B
B_cells
CD4_T_cells
CD8_T_cells
CD8_T_cells_PD1_high
CD8_T_cells_PD1_low
Endothelium
Fibroblasts
Lymphocytes
Macrophages
Macrophages_M1
Macrophages_M2
Monocytes
NK_cells
Neutrophils
Non_plasma_B_cells
Plasma_B_cells
T_cells
T_helpers
Tregs
microbes number
C
Chitinivibrio
Endothelium
Campylobacter
Macrophages_M2
Macrophages
CD8_T_cells
Tregs
Flammeovirga
CD8_T_cells_PD1_high
CD8_T_cells_PD1_low
Lachnoclostridium
Macrophages_M1
T_cells
Lymphocytes
NK_cells
CD4_T_cells
Collimonas
Fibroblasts
T_helpers
Plasma_B_cells
Terrabacter
Sutterella
B_cells
edges
immune cells
microbes
Endothelium
Fibroblast
rho

**Figure S2. Sample size in this study and the associations between tumor microbes and cell types. A)** Bar plot showing the number of samples with only eukaryota (blue), only bacteria/viruses/archaea (green), or both (yellow) by cancer type (x-axis). **B-C)** The distribution and the network of the associations between pan-cancer key microbes and cell types in 32 cancer types. The size of microbe nodes represents size of node. Edges in **C** weighted by the Spearman's correlation coefficients.

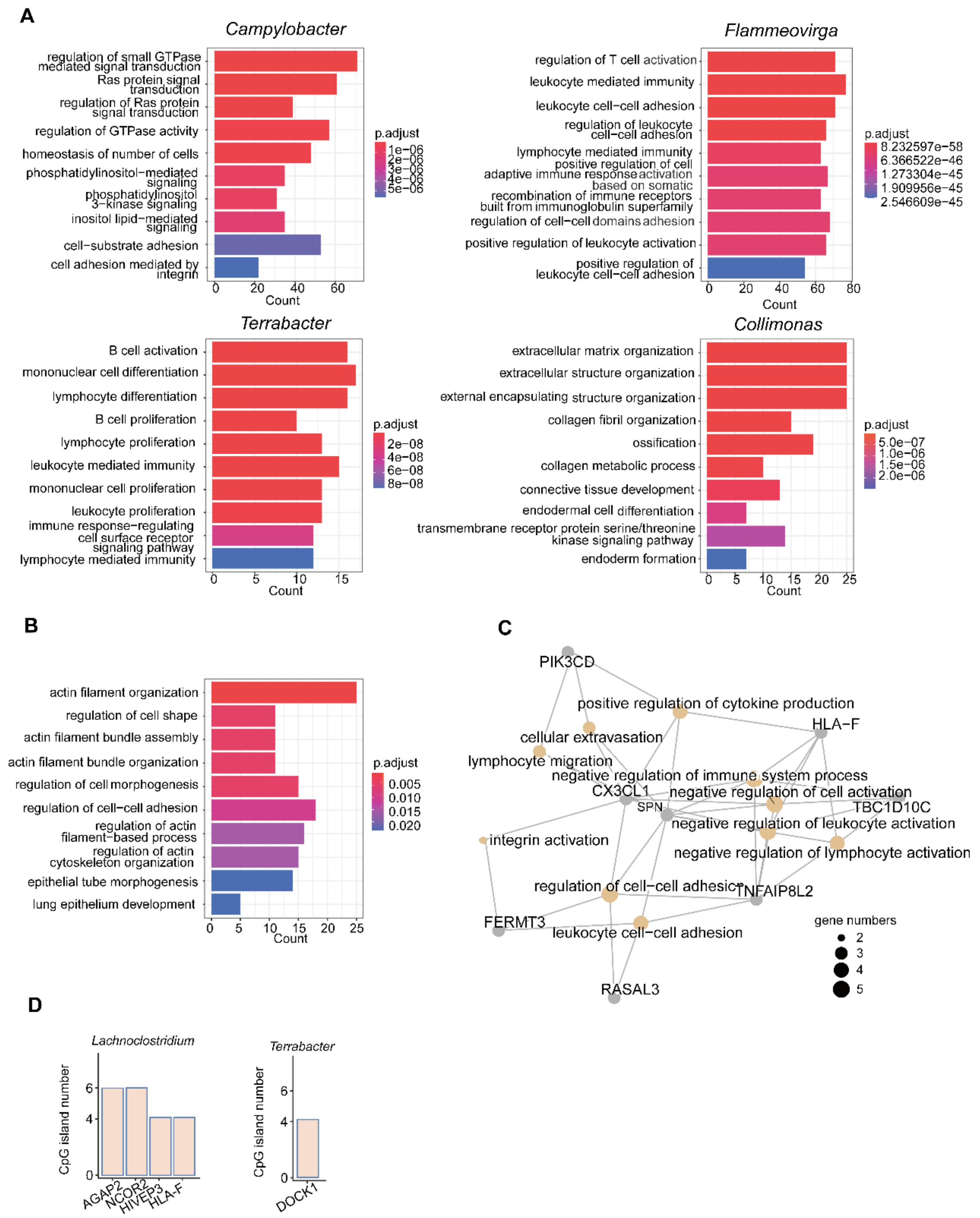


**Figure S3. GO enrichment analysis of genes associated with pan-cancer key tumor microbes. A)** GO enrichment analysis of genes associated with four pan-cancer immune-related microbes (*Campylobacter*, *Flammeovirga*, *Terrabacter*, *Collimonas*). Top 10 pathways are showed here based on adjusted *p* values. **B**) GO enrichment analysis of genes linked to CpG islands associated with immune-related microbes (*Campylobacter*, *Flammeovirga*, *Terrabacter*, *Collimonas*) **C)** GO enrichment analysis of genes linked to CpG islands associated with *Lachnoclostridium*, and top 10 pathways are showed here based on gene ratio. The right panel shows genes linked to multiple CpG islands. **D**) Multiple associated CpG islands in single gene of *Lachnoclostridium*.

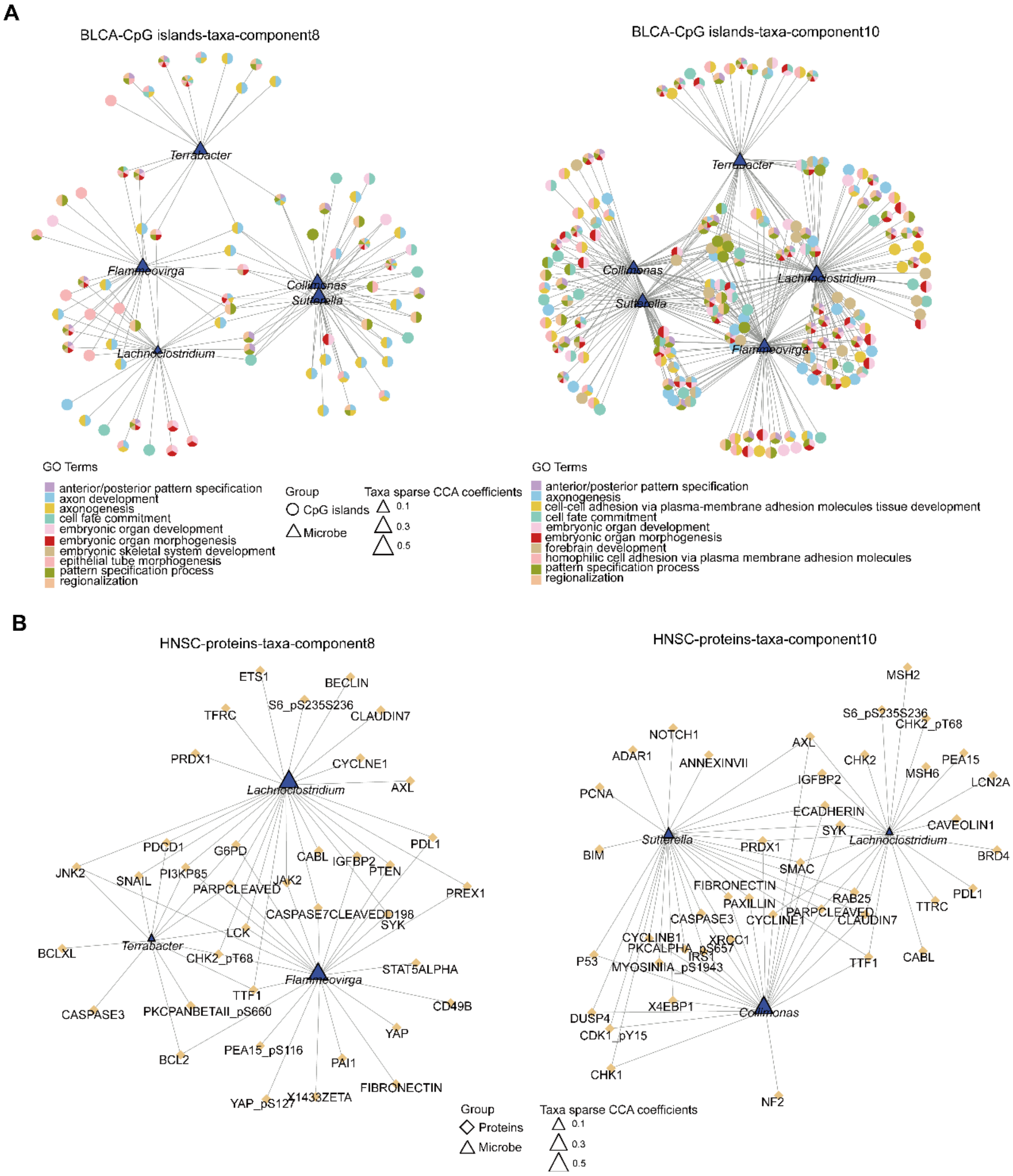


**Figure S4. Differential contribution of cluster 1 and cluster 2 pan-cancer microbes in different host pathways at the DNA methylation and protein expression levels** **A)** Host GO pathways enriched in genes related to CpG islands associated with pan-cancer key microbes in the component 8 and 10 of BLCA based on sparse CCA. Dot represents gene and its GO terms, and triangle size represents the absolute value of sparse CCA coefficients for microbes. Among each component, only host-microbe associations with Spearman's rho > 0.3 were used. Only top 10 GO terms based on adjust *p* values are showed here. **B)** Proteins associated with pan-cancer key microbes in the component 8 and 10 of HNSC based on sparse CCA. Dot represents proteins, and triangle size represents the absolute value of sparse CCA coefficients of microbes. Among each component, only associations with Spearman's rho > 0.3 were kept. Only top 10 GO terms based on adjust *p* values are showed.

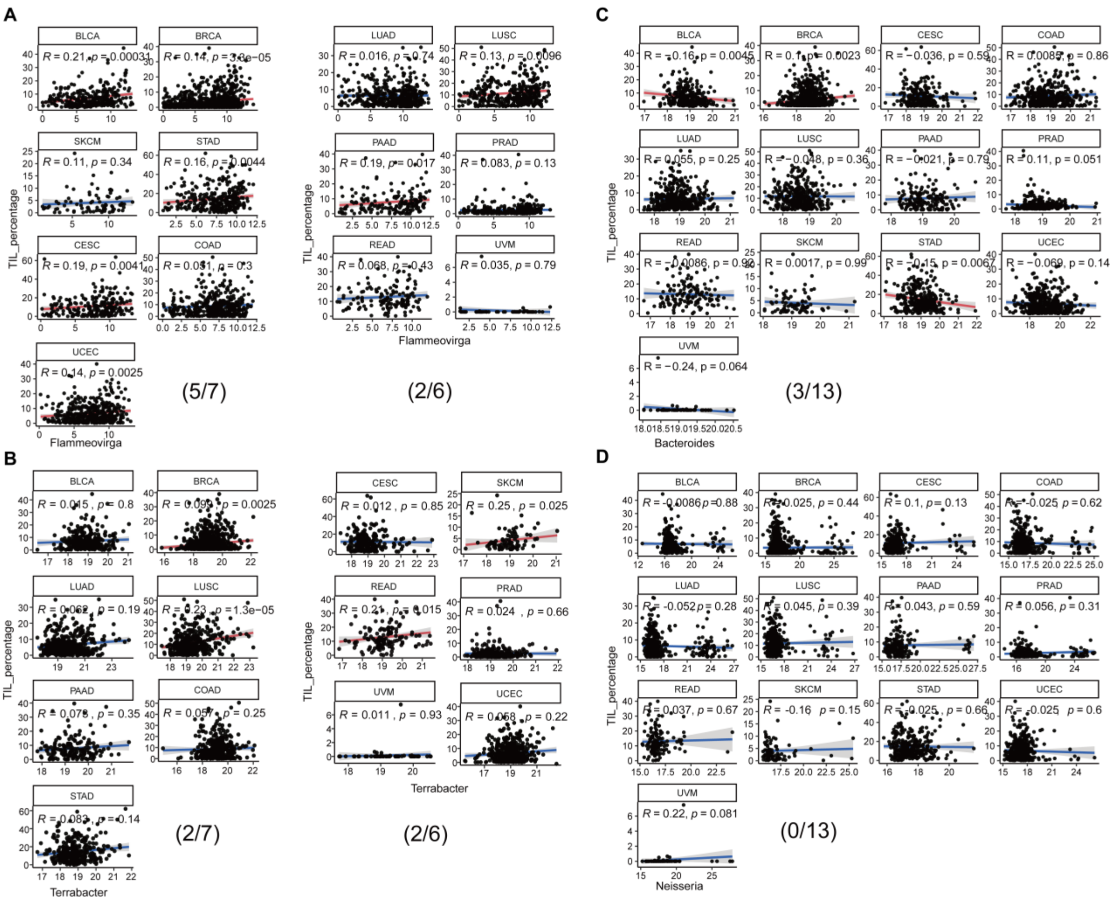


**Figure S5. The associations between pan-cancer key microbes and TIL (tumor-infiltrating lymphocytes) percentage. A)** *Flammeovirga*. **B)** *Terrabacter*. **C)** Control *Bacteroides*. **D)** Control *Neisseria*. For each panel, the left shows cancer types where microbes were associated lymphocytes in the initial test using cell type profiles from RNA-Seq deconvolution, while the right shows the rest cancer types with TIL percentage data. The red line represents significant association (Spearman's rho test $p < 0.05$), and the blue line represents no significant correlation. The number within the parentheses of each panel (such as 5/7 and 2/6 in panel A) shows the number of significant cancer types over the total number of cancer types.

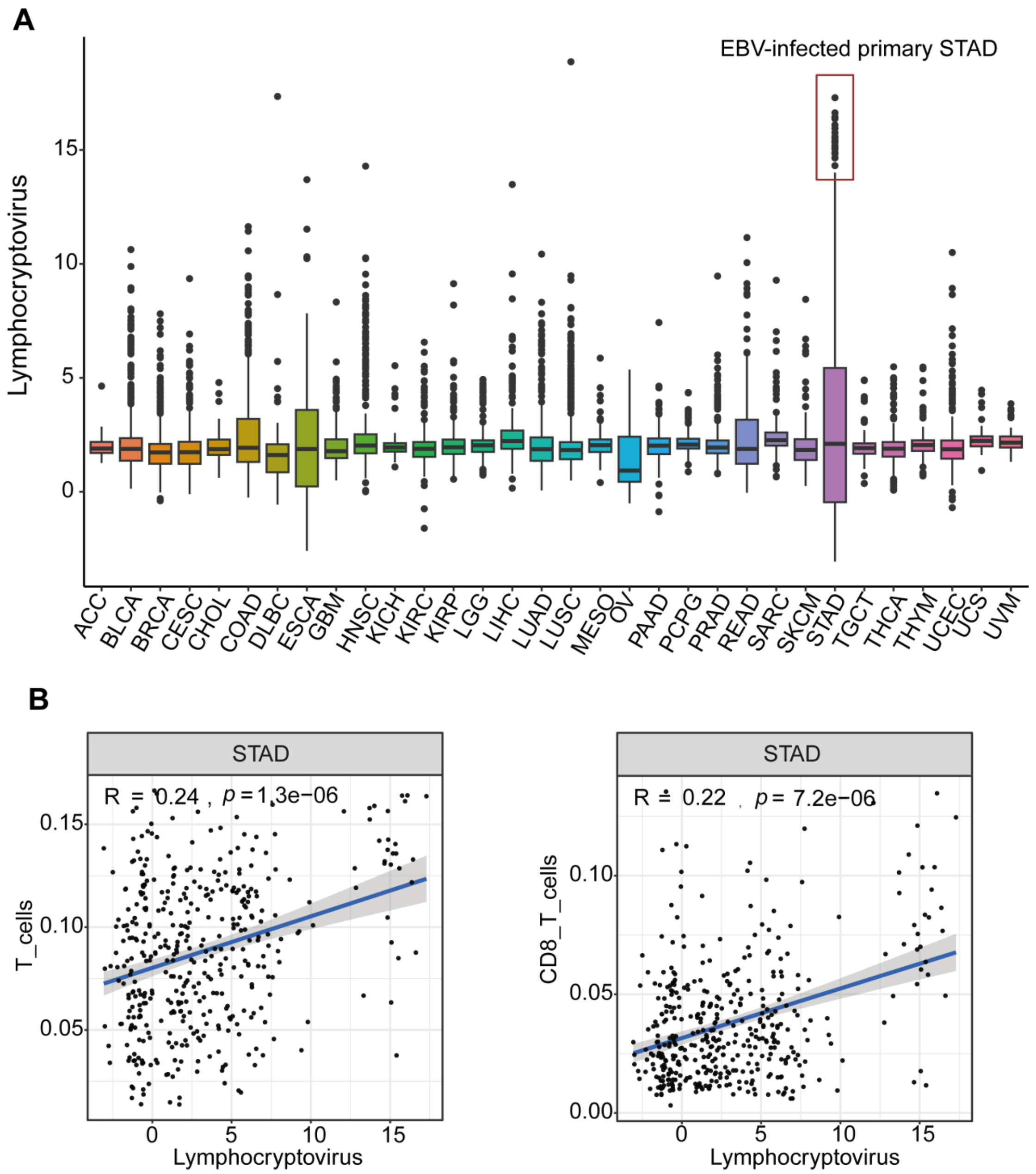


**Figure S6. The abundance of *Lymphocryptovirus* in 32 cancer types and its associations with T cells. A)** The abundance of *Lymphocryptovirus* (EBV) in 32 cancer types, with EBV–infected samples highlighted. **B)** The associations between the abundance of *Lymphocryptovirus* with T cells and CD8+ T cells. Spearman's rho and *p* values are showed.

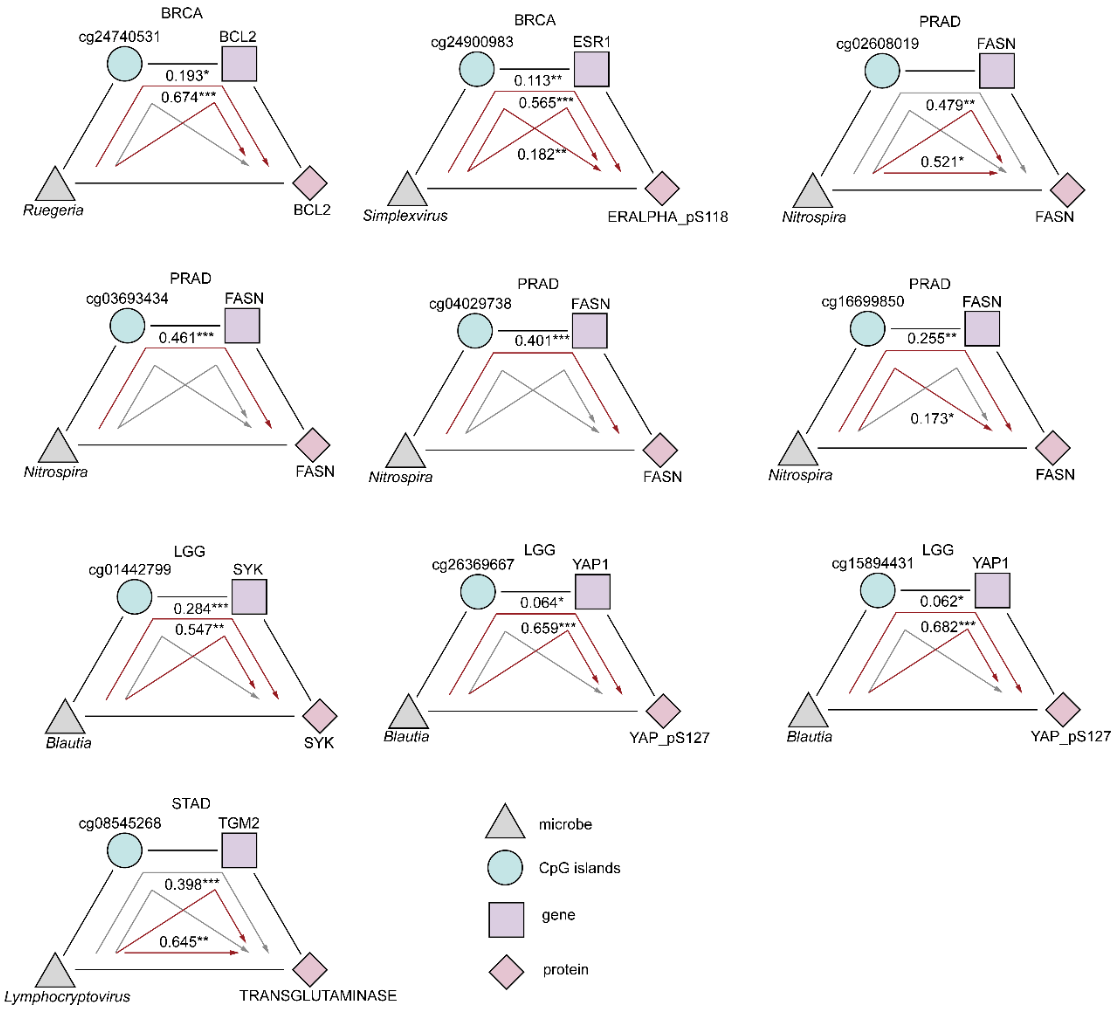


**Figure S7. Mediation analysis between tumor microbes and host molecular features.** Each plot represents a potential microbe-methylation-RNA-protein axis from Figure 4D. The proportional effect size (shown in number) and p value (*, <0.05; **, <0.01; ***, <0.001) are showed in each plot, and the red line represents significant path with $p < 0.05$ and non-zero effect size.

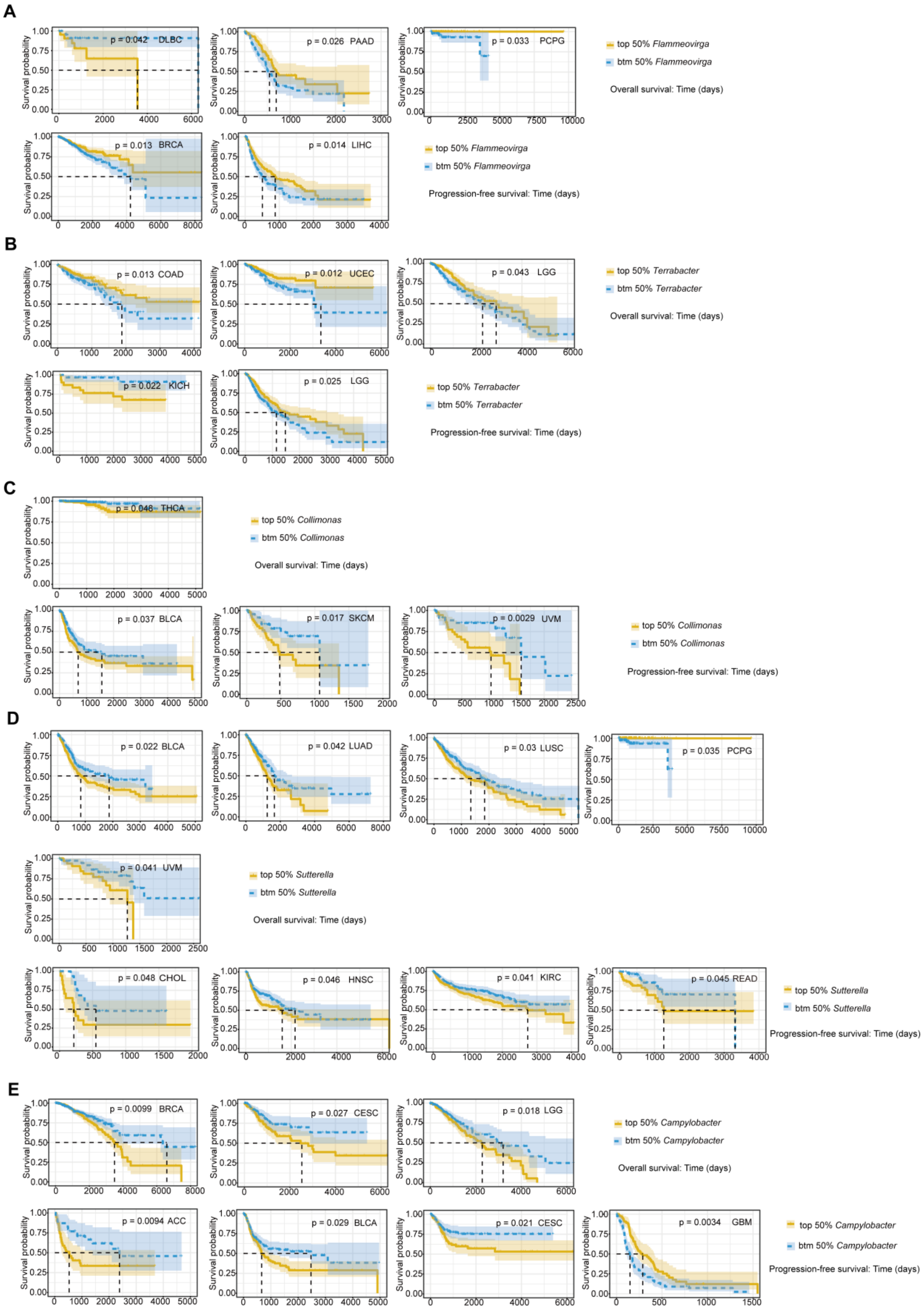


**Figure S8. Survival analysis of pan-cancer key microbes. A)** *Flammeovirga*. **B)** *Terrabacter*. **C)** *Collimona*. **D)** *Sutterella*. **E)** *Campylobacter*. Blue represents top 50% samples, while orange represents bottom 50% samples based on the abundance of microbes. Results with $p < 0.05$ are showed.
